# VieMedKG: Knowledge Graph and Benchmark for Traditional Vietnamese Medicine

**DOI:** 10.1101/2024.08.07.606195

**Authors:** Tam Trinh, Anh Dao, Hy Thi Hong Nhung, Hy Truong Son

## Abstract

Traditional Vietnamese Medicine (TVM) and Traditional Chinese Medicine (TCM) have shared significant similarities due to their geographical location, cultural exchanges, and hot and humid climatic conditions. However, unlike TCM, which has substantial works published to construct a knowledge graph, there is a notable absence of a comprehensive knowledge graph for TVM. This paper presents the first endeavor to build a knowledge graph for TVM based on extensive existing resources from TCM. We name our knowledge graph as VieMedKG. We propose a translation and filtration process to adapt TCM knowledge graphs to TVM, identifying the overlapping and unique elements of TVM. In addition, the constructed knowledge graph is then exploited further for developing a curated benchmark for the knowledge graph-based question-answering problem with the potential to support doctors and patients in assisting doctors and patients in identifying various diseases. Our work will not only bridge the gap between TCM and TVM but also set the foundation for future research into traditional Vietnamese medicine community. Our source code is publicly available at https://github.com/HySonLab/VieMedKG/.

## 1 Introduction

Traditional Vietnamese Medicine (TVM) is a complete medical system that has been discovered and developed for many thousands of years. In the past, deeply influenced by the ancient philosophies of Yin and Yang, the Five Elements from Traditional Chinese Medicine, TVM has integrated a wide range of elements, including herbal medicine, massage, disease names, and methods to cure them, while also preserving and integrating the unique Vietnamese traditional elements and indigenous knowledge. Since TVM has evolved throughout the long history of Vietnamese nation and people, a systematic way such as Knowledge Graph to represent a comprehensive view of TVM is crucial. A knowledge graph can effectively organize and represent the complex relationship between various element of TVM such as diseases, symptoms, herbal medicine, the cause of disease and treatment methods. This structured representation can not only help cultural preservation but also make TVM more accessible to develop future research works and applications in the domain of Vietnamese traditional medicine.

However, despite several advantages, building a knowledge graph from a vast resources of TVM requires overcoming many challenges. The knowledge of TVM is often documented in historical texts, local manuscrips or oral communication which are inherently unstructured and lack of standardardiztion. Knowledge Graphs help to prevent hallucination in Large Language Models (LLMs) because they are more rigorous and structured. Therefore, processing these kind of data into a knowledge graph for TVM can be time consuming and costly. At the same time, many works (Chen et al., 2023; Cheng et al., 2018; Long et al., 2019; Zhou et al., 2010) have been published with significant efforts to collect, validate, contruct and exploit several downstream application of TCM in knowledge graph representation such as graphbased question answering and infomation retrieval. Realizing the similarities between TCM and TVM, we propose a method to leverage the existing TCM knowledge graph as a foundation in order to build a comprehensive knowledge graph for TVM. Our approach involves not only translating TCM data into Vietnamses version but also filtering and adapting that data to preverse and reflect the uniqueness of Vietnamese traditional medicine. By doing so, we can effectively utilize the huge amount of TCM data already available. Furthermore, we also analysis and propose a benchmark for graph-based question answering problem on our TVM knowledge graph. To sum up, our contributions can be divided into the following categories:

- We introduce a new knowledge graph VieMedKG for Traditional Vietnamese Medicine built upon the current existing Traditional Chinese Medicine by applying translation and filtration process. To the best of our knowledge, this is the first endeavor to construct a systemetic way to represent TVM knowledge (see Figure 1).
- We propose a curated benchmark for graph-based question answering problem about TVM and further analysis some methods on VieMedKG.

**Figure 1.**
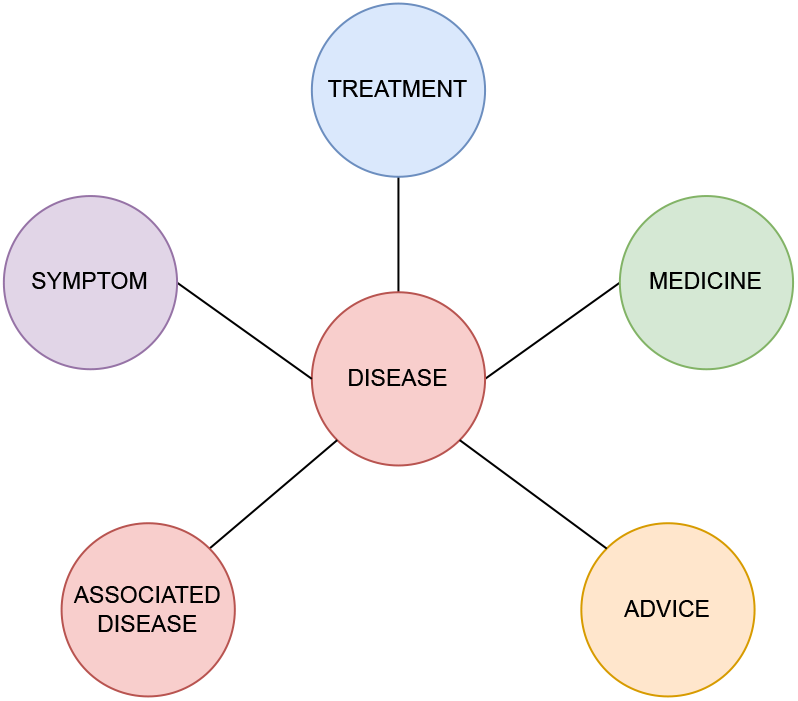
Illustrative representation of VieMedKG, showing the connections among entities.

## 2 Related works

### 2.1 TCM and TVM dataset

The availability of comprehensive datasets is crucial for knowledge discovery in TCM. Several significant TCM data resources have been developed. China TCM Patent Database (CTCMPD), managed by the State Intellectual Property Office of China, includes over 19,000 patents and 40,000 formulae (Liu and Sun, 2004). TCMBank (Lv et al., 2023), extended from TCM Database@Taiwan (Chen, 2011), includes 9,192 herbs, 61,966 ingredients, 15,179 targets, and 32,529 diseases.

In addition, several notable efforts have been made to construct TCM Knowledge Graphs. Yu et al. (2017) constructed the TCM health care knowledge graph through ontology-based database integration. Zheng et al. (2020) constructed TCMKG by training deep learning model for named entity recognition, aligning data, and creating triples based on the ontology. Zhang and Hao (2024) adopted Large Language Models for name entity recognition to construct a knowledge graph of TCM.

In the context of Vietnamese Medicine, several datasets have been developed to facilitate research and application in this field. VietHERB, a database for Vietnamese Herbal Species, was built in an ontology-based structure to organize the numerous hierarchical multi-class of herbs (Nguyen-Vo et al., 2018). However, Nguyen-Vo et al. (2018) pointed out that the limitations of obtaining data are poor handwriting records and unexpected incorrect information when computerizing from hard-copied documents. ViMQ, a Vietnamese Medical Question Dataset, contains 9,000 questions for the two tasks of intent classification and named entity recognition (Huy et al., 2021). Despite these resources, there is no benchmark for Question Answering (QA) over Knowledge Graphs (KGs) for Vietnamese Medicine, highlighting a gap that needs to be addressed for further advancement in this field.

### 2.2 Question Answering System using Knowledge Graphs (KGs)

Early efforts in Knowledge Graph Question Answering (KGQA) aimed to convert natural language queries into formal logical queries using deep neural semantic parsing such as (Andreas et al., 2016; Lan et al., 2019; Yin et al., 2016). However, these approaches struggle with complex queries that need multi-hop reasoning and managing constraints on incomplete knowledge graphs. An alternative research approach to Knowledge Graph Question Answering (KGQA) frames the task as a graph traversal problem, directly reasoning on the input graph to uncover structural patterns in transitive relations. Methods like VRN (Zhang et al., 2018), GraftNet (Sun et al., 2018), and PullNet (Sun et al., 2019) employ graph neural networks and local subgraph searches to navigate entities within the knowledge graph.

Recent advancements in integrating large language models (LLMs) with knowledge graphs (KGs) have shown significant progress. Techniques such as REALM (Guu et al., 2020) and RAG (Lewis et al., 2020) have been developed to retrieve documents and enhance language models using these documents. Furthermore, KGs can serve as a valuable & rigorous knowledge source, where information is succinctly encoded, and several methods have been proposed to integrate these facts from KGs into language models (Galetzka et al., 2021; Rony et al., 2022; Kang et al., 2022). Nevertheless, these approaches generally require extensive training data and frequent model updates for various downstream tasks. Recent research (Izacard et al., 2024) indicates that retrieval-augmented language models can achieve strong performance with few-shot learning. In our work, we will experiment with retrieval-augmented language models using both zero-shot and few-shot learning, without additional training steps.

## 3 Method

### 3.1 Knowledge Graph for Traditional Vietnamese Medicine (VieMedKG)

#### 3.1.1 Access to Traditional Medical Knowledge

Both TVM and TCM share a historical foundation due to Vietnam’s annexation by China for over a thousand years, facilitating the transfer of medical knowledge and practices. Both traditions heavily rely on herbal remedies, but TVM adapts its pharmacopeia to include local Vietnamese plants suited to the tropical climate, unlike the temperate-climate plants used in Chinese Medicine (Bui, 2019). Additionally, TVM data collection faces challenges like poor handwriting in historical records and errors during computerization (Nguyen-Vo et al., 2018). In contrast, TCM benefits from a well-established and comprehensive database developed over several decades. As a result, leveraging the TCM database can improve the accuracy and depth of medical research. In this study, we obtained structured data from OpenKG, the Chinese open database. Our dataset, presented in Traditional Chinese, contains comprehensive information on 9,464 different diseases, including their symptoms, treatments, medications, patient advice, and linked diseases.

#### 3.1.2 Translation

Vietnamese word formation has been significantly influenced by Chinese, with Chinese-originated words accounting for about 70% of Vietnamese vocabulary, termed Sino-Vietnamese, and about 50% in modern Vietnamese (Ðăng, 2003). Sino-Vietnamese can be easily translated word by word, often resulting in meaningless translations, and some terms lack Vietnamese equivalents but have English ones. Therefore, we have decided to translate from Chinese to English, then from English to Vietnamese.

Baidu Translator is widely recognized as the most popular tool for translating Chinese to English quickly through free internet use (Yao et al., 2012). According to (Razzaka et al., 2019), Baidu translation tends to be more accurate and contextually appropriate for traditional Chinese literature than Google Translator, which is one of the most popular translators. Furthermore, Baidu Translator offers a specialized API for medical translations. For instance, the Chinese term “附錄” is translated by Google Translator as “ph ụ lục,” referring to a section at the end of a book, whereas Baidu Translator renders it as “ruôt thùa,” aligning with its medical meaning as a small organ attached to the intestines on the right side of the human body.

Consequently, we have chosen to utilize Baidu Translator for translation. To ensure accuracy and the quality of the translations, we then used GPT-4 (Achiam et al., 2023) and human evaluation, which is checked by a linguistic expert with a PhD in Chinese linguistics for fact-checking, translating quality, and refinement.

#### 3.1.3 Construction of VieMedKG

The construction of VietMedKG requires a number of processes such as Entity Disambiguation and Entity Linking. The core of the knowledge graph are the triple’s entities, attributes, and relationships. The entities and attributes are combined and fed into the knowledge graph through the popular graph database - Neo4j database (Miller, 2013) to store data about nodes and relationships. Entities have 5 labels including disease, symptom, treatment, medicine, and advice corresponding to which relationships include HAS_SYMPTOM, HAS_TREATMENT, USE_MEDICINE, HAS_ ADVICE, and IS_LINKED_WITH. Each entity provides its own additional attributes with detailed information as Figure 4 shows. The constructed medical data knowledge graph includes 41,549 nodes and 42,112 relationships. The detailed description of the knowledge graph is shown in Table 3 and Table 4.

**Figure 2.**
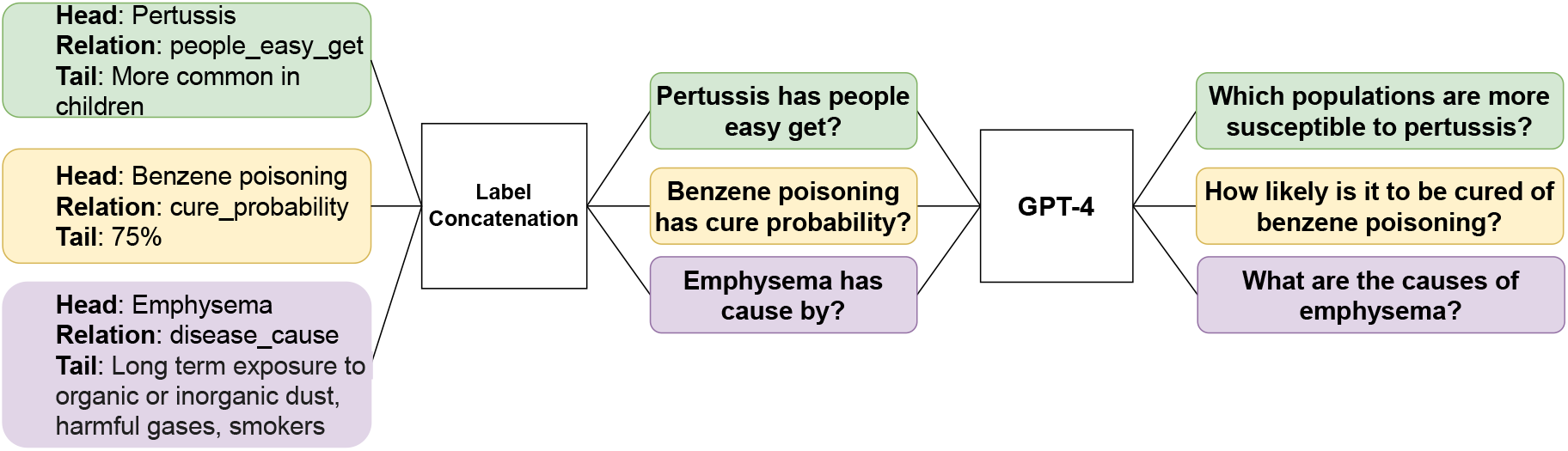
Overview of the procedure for question generation. The input is a triple from the KG, and the output is a natural language question.

**Figure 3.**
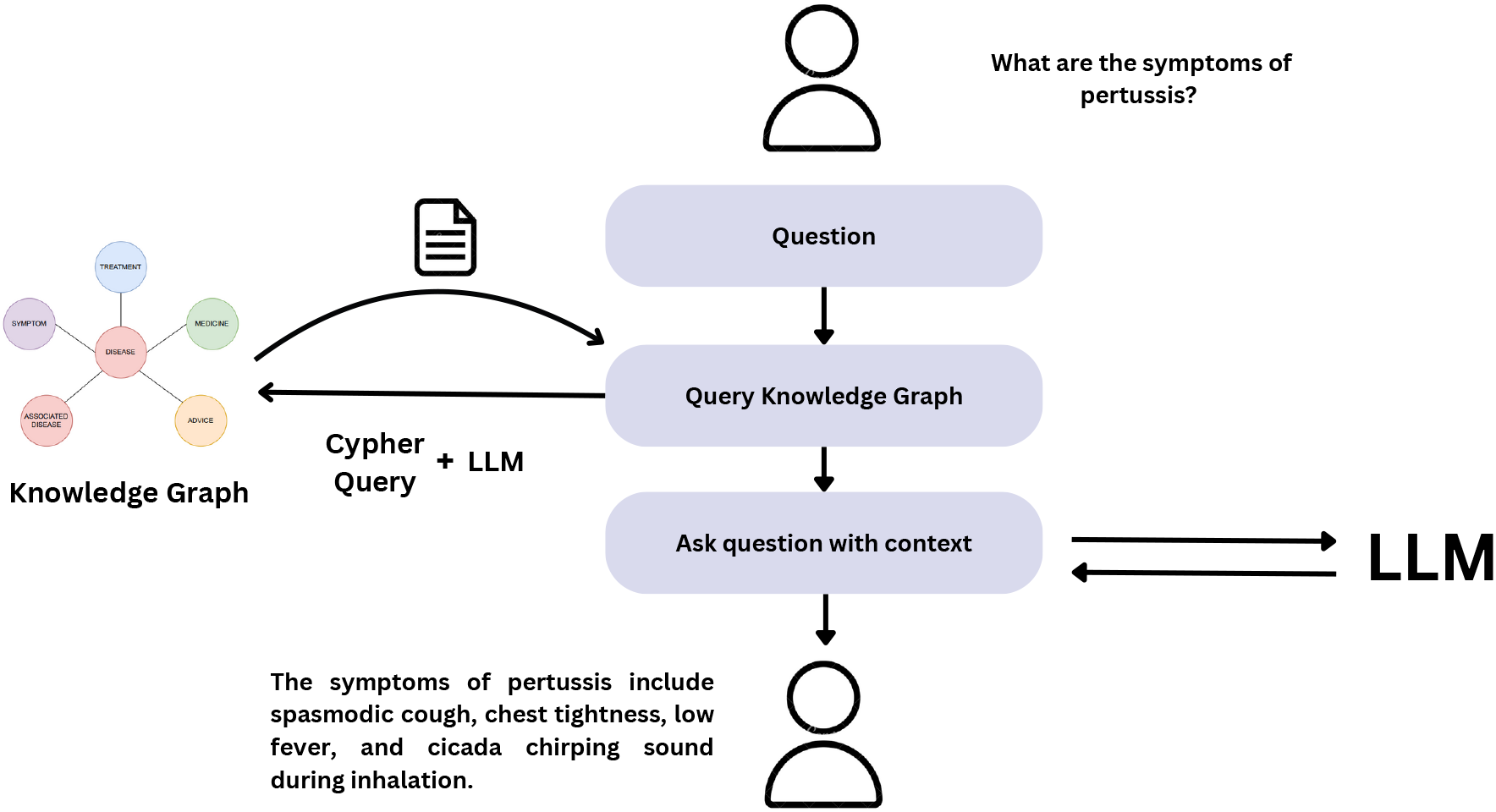
Overview of the RAG system. The process begins with a user question processed through Cypher and LLM queries to retrieve relevant knowledge graph nodes. An answer is then generated using the augmented query and contextual information.

**Figure 4.**
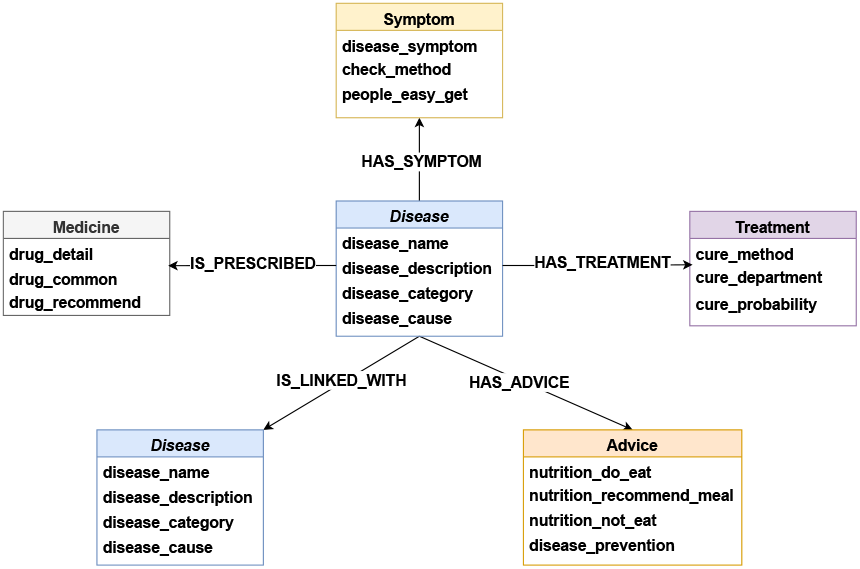
Graph Schema for VietMedKG.

**Figure 5.**
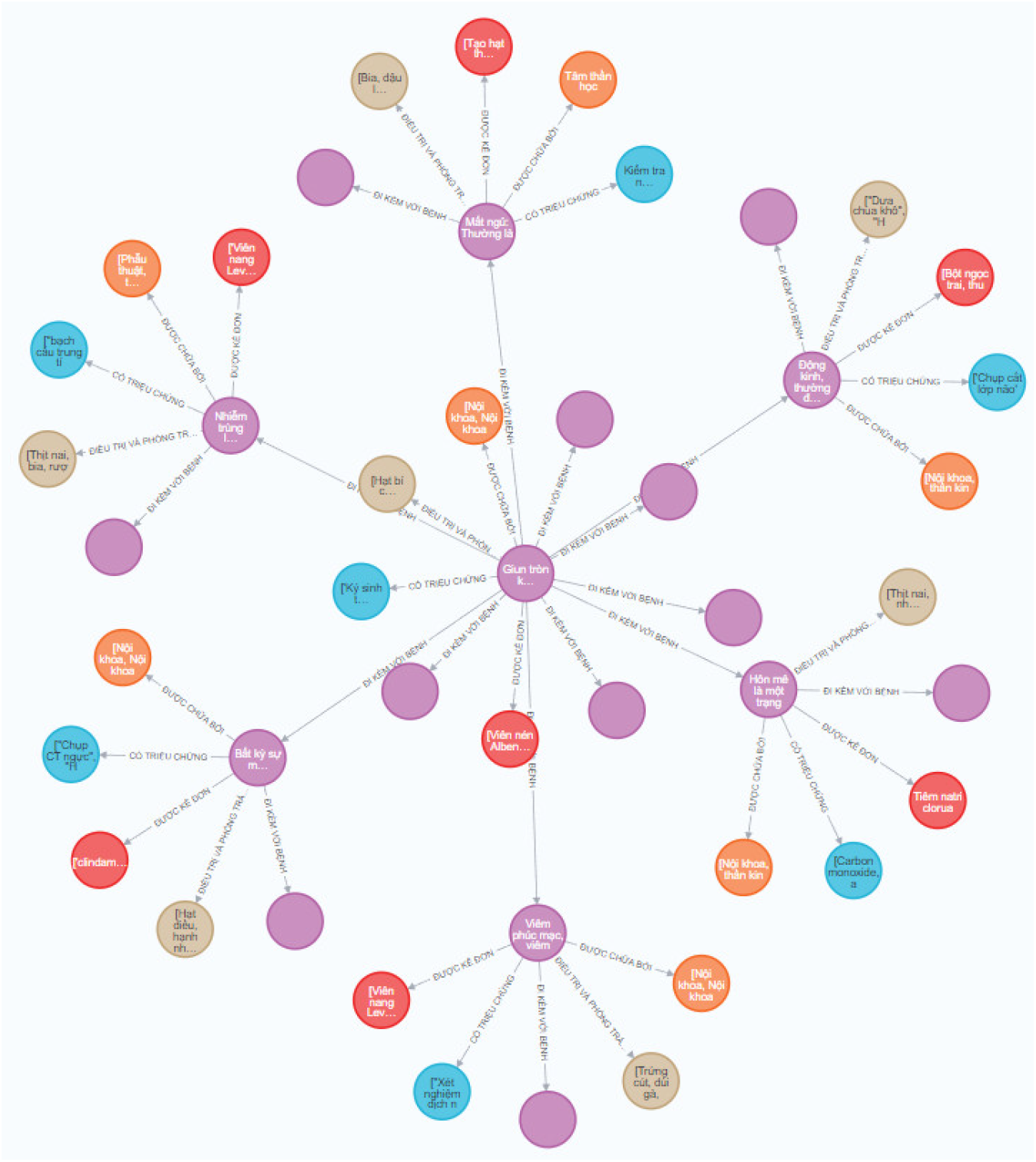
Subgraph of VieMedKG, visualized using Neo4j.

**Table 1:** Results for 1-hop and 2-hop retrieval performance.

| Model | 1-hop | 2-hop |
| --- | --- | --- |
| GPT4 | 0.4143 | 0.4 |
| Gemini 1.0 Pro | 0.5075 | 0.4909 |

**Table 2:** Results for 1-hop and 2-hop question-answering performance. We highlight the best result for each metric.

| Model | 1-hop |  |  | 2-hop |  |  |
| --- | --- | --- | --- | --- | --- | --- |
|  | BLEU | METEOR | ROUGE | BLEU | METEOR | ROUGE |
| GPT4 | 0.058 | 0.157 | 0.104 | 0.021 | 0.092 | 0.112 |
| Gemini 1.0 Pro | 0.051 | 0.14 | 0.123 | 0.035 | 0.126 | 0.118 |
| GPT4 + RAG (ours) | **0.417** | **0.552** | **0.508** | **0.235** | **0.325** | **0.298** |
| Gemini 1.0 Pro + RAG (ours) | 0.345 | 0.492 | 0.479 | 0.204 | 0.265 | 0.246 |

**Table 3:** Number of Entities by Type.

| Entity Type | No. of Entities |
| --- | --- |
| disease | 9,464 |
| symptom | 8,806 |
| treatment | 8,806 |
| medicine | 8,806 |
| advice | 5,577 |
| **Total** | **41,549** |

**Table 4:** Number of Relationships by Type.

| Relationship Type | No. of Relationships |
| --- | --- |
| HAS_SYMPTOM | 8,806 |
| HAS_TREATMENT | 8,806 |
| USE_MEDICINE | 8,806 |
| HAS_ADVICE | 5,577 |
| IS_LINKED_WITH | 10,117 |
| **Total** | **42,112** |

VieMedKG is stored in Neo4j Graph Database, a NoSQL database that falls under the category of graph databases, adhering to the mathematical theory of trees. In Neo4j, nodes are represented as vertices and relationships as edges. A literature review by (López and la Cruz, 2015), indicates that Neo4j demonstrates superior performance in terms of time response and provides shorter pathfinding compared to relational databases.

#### 3.1.4 Benchmark Dataset for Question Answering

##### Triple creation

In constructing a benchmark dataset for question-answering systems, the process of triple creation is pivotal. A triple is composed of three components: a head, a relation, and a tail. In the context of a dataset centered on diseases, the head represents the name of the disease, the relation denotes the attribute associated with the disease, and the tail indicates the value of that attribute. These triples are then used to generate various types of questions, including 1-hop and 2-hop questions, which assess the ability of question-answering systems to navigate different levels of complexity in the knowledge base.

We used the two sequential approaches which are label concatenation method and the usage of Large Language Model (LLM). The initial method involves concatenating labels from unseen triples to generate questions. For instance, using the triple (“*pertussis*”, “*people_easy_get*”, “*More common in children*”) would create a question such as “*Pertussis have people easy get?*”. Despite potentially generating grammatically incorrect or nonsensical questions, this approach effectively imparts textual information to the model concerning keywords linked with the relations.

**1-hop question dataset** For 1-hop question, we designed 32 types of questions in total (see Table 5). In exploring the relationship between extensive medical information (an attribute in the graph) and disease identification, this study encountered challenges with question length. Utilizing all details from attributes resulted in overly long questions. To address this, the GPT-4 language model (Achiam et al., 2023) was employed. GPT-4 offered two functionalities: 1) random summarization of an attribute, and 2) targeted selection of a key point within that attribute. Both approaches aimed to shorten the question to under 25 words while preserving its core meaning.

**Table 5:**
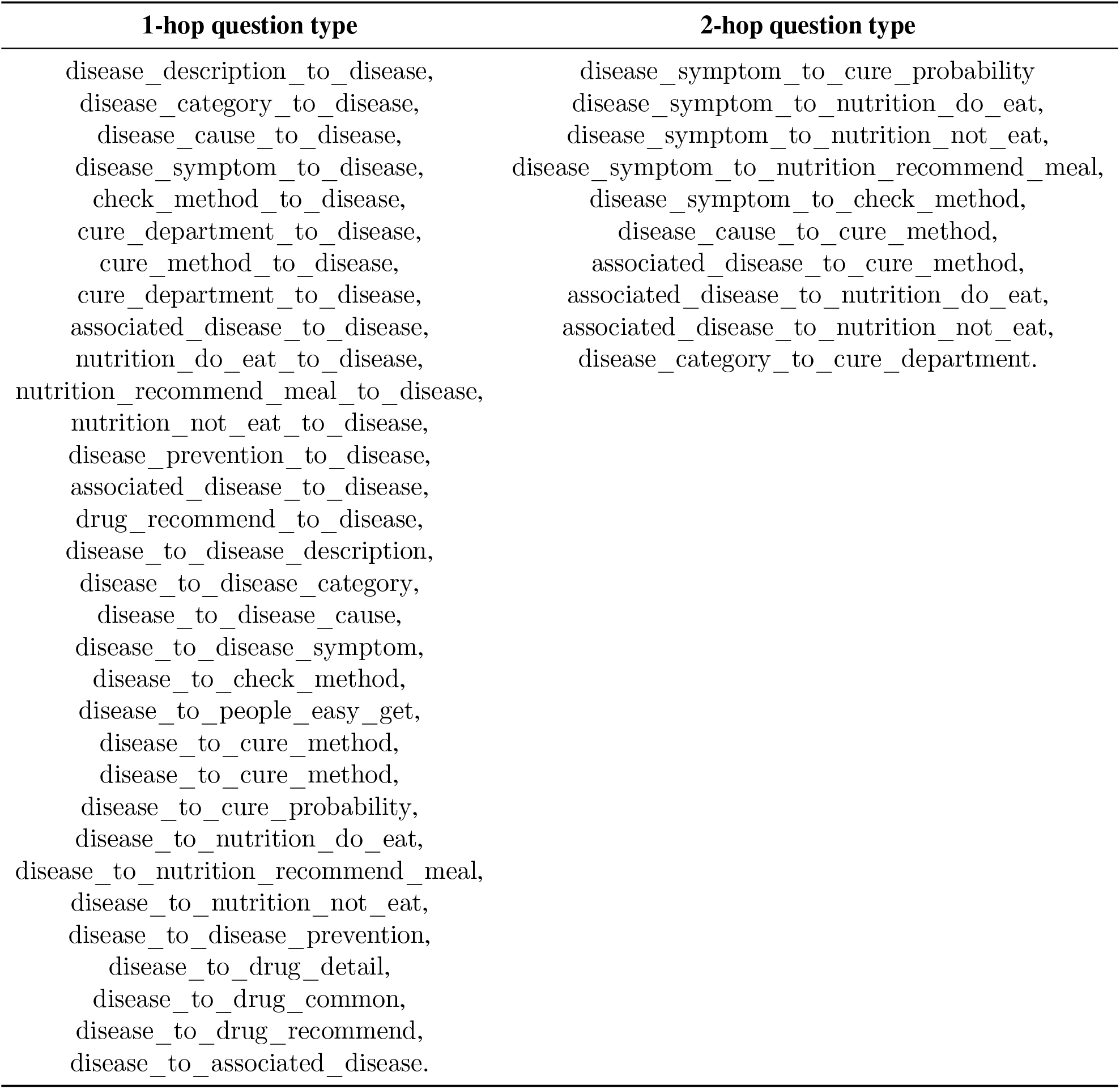
Question types of 1-hop and 2-hop questions.

**2-hop question dataset** We designed 10 types of questions (see Table 5). To address the issue of lengthy questions, we employed a similar methodology as used for the 1-hop questions.

### Creating multiple answer questions

We grouped questions of the same type from the dataset and calculated the Jaccard similarity (Niwattanakul et al., 2013) between attribute values used in question creation. If the similarity exceeded 0.9, we merged the answers accordingly. This process ensured that questions with highly similar attributes yielded merged answers, optimizing question diversity while maintaining relevance.

### Generating natural-sounding question using LLM

We employed the GPT-4 language model to create natural-sounding questions from those generated by the label concatenation method. Using zero-shot prompting, we instructed the model to emulate doctors or patients seeking disease information. Figure 2 demonstrates the question generation process from triples, involving feeding pairs of triples and target questions derived from the training data.

By applying GPT-4 (Achiam et al., 2023) to our training set, we harnessed its ability to encode relationships between words and phrases, thus producing coherent questions even for triples with previously unseen relations. Subsequently, we manually reviewed and eliminated nonsensical questions, and utilized uniform sampling to establish the benchmark. The dataset consists of 4,232 questions, including 2,164 one-hop questions and 2,068 two-hop questions. Table 6 provides a detailed description of the benchmark.

**Table 6:**
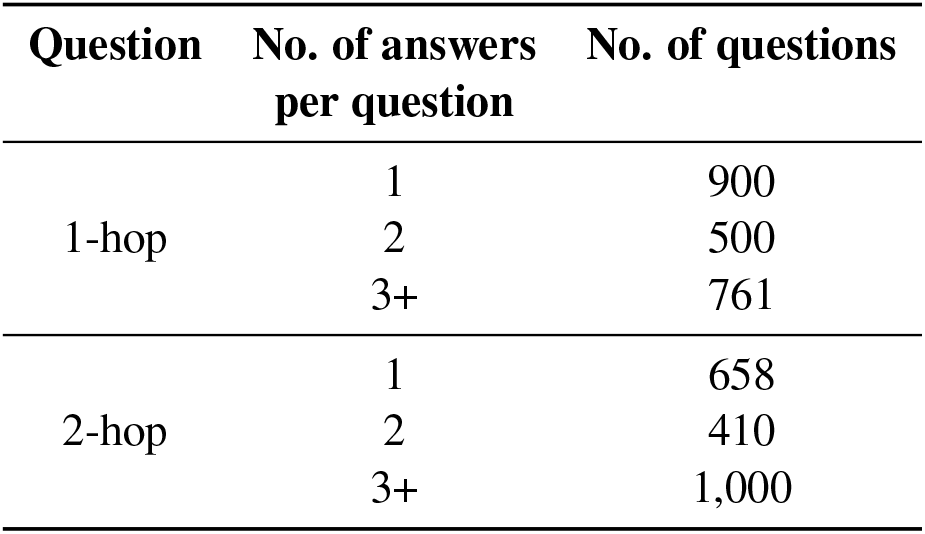
Description of KGQA benchmark.

| Question | No. of answers<br>per question | No. of questions |
| --- | --- | --- |
| 1-hop | 1 | 900 |
|  | 2 | 500 |
|  | 3+ | 761 |
| 2-hop | 1 | 658 |
|  | 2 | 410 |
|  | 3+ | 1,000 |

### 3.2 Retrieval-Augmented Generation (RAG) with VietMedKG

#### 3.2.1 The Cypher Query

Neo4j’s query language is called Cypher (Francis et al., 2018). Cypher is a declarative query language for graphs that uses graph pattern-matching as a main mechanism for graph data selection for both read-only and mutating operations. As an example, the query to retrieve symptoms of pertussis can be stated as follows:

START Disease=node:

Disease(disease_name={disease_name})

MATCH (Disease)-[:HAS_SYMPTOM] ->

(Symptom:disease_symptom)

WHERE Sisease.disease_name = ‘pertussis’

RETURN Symptom.disease_symptom, COUNT(*)

ORDER BY COUNT(*) DESC

This query starts by searching the index called “Disease” for a node whose attribute “disease_name” is set to the contents of the parameter “disease_name”. It then uses pattern matching to find all symptoms associated with this disease. The “WHERE” clause ensures that we are specifically querying for the disease “pertussis”. As a result, the query returns all symptoms associated with pertussis and how often they were encountered during the traversal, which is also the sort criterion. Attributes could be accessed by “Symptom.disease_symptom”.

To enhance the effectiveness of Cypher statements in practice, leveraging the few-shot learning capabilities of large language models (LLMs) involves presenting examples that guide the model in generating Cypher statements. This approach facilitates the adaptation of Cypher syntax to specific query requirements, ensuring compatibility with database structures and query outcomes.

#### 3.2.2 Augmenting LLM answer with RAG and zero-shot prompting

Figure 3 illustrates the architecture of the Retrieval-Augmented Generation (RAG) system. The workflow begins with the user submitting a question. This question is then processed using a Cypher query and LLM to generate relevant queries. The retrieved nodes are used to query the knowledge graph through Cypher queries, extracting additional contextual information. The results are returned in JSON format, after which LLM generates an answer based on the augmented query and the contextual information provided by the knowledge graph.

## 4 Experiments

### 4.1 Experiment Design

Researchers have been working on Question-Answering (QA) system for Vietnamese such as (Tran et al., 2023). However, in this paper, we propose the first-ever QA system that incorporates Knowledge Graph (i.e. VieMedKG) for medicine in Vietnamese. We evaluated our QA system using a benchmark dataset designed for QA tasks based on VieMedKG. In our experiments, we employed large language models (LLMs), specifically GPT-4 (Achiam et al., 2023) and Gemini 1.0 Pro (Team et al., 2023), using zero-shot prompting to serve as baseline for the benchmark. Subsequently, we leveraged our RAG system with the GPT-4 and Gemini 1.0 Pro as the generator to measure the performance for both 1-hop and 2-hop benchmark dataset.

To evaluate the retrieval performance and determine the contribution of retrieved triples to answer generation, we employed accuracy as our primary metric. For assessing the overall performance of the QA system, we utilized metrics including BLEU (Papineni et al., 2002), ROUGE (Lin, 2004), and METEOR (Lavie and Agarwal, 2007) scores.

### 4.2 Experimental Results and Analyses

The performances of retrieval and questionanswering are detailed in Tables 1 and 2, respectively. When compared to the zero-shot prompting capabilities of state-of-the-art language models such as GPT-4 and Gemini 1.0 Pro, our system exhibits a significant improvement in questionanswering performance. Specifically, our proposed RAG system enhances the BLEU score of GPT-4 by approximately 7.19 times and that of Gemini 1.0 Pro by approximately 6.76 times for 1-hop questions, and by approximately 11.19 times for GPT-4 and 5.83 times for Gemini 1.0 Pro for 2-hop questions. These considerable improvements come with only a marginal increase in inference latency, adding an average of 8.40 seconds for GPT-4 and 5.52 seconds for Gemini 1.0 Pro.

## 5 Conclusion

In this paper, we have introduced the first knowledge graph for Traditional Vietnamese Medicine (TVM), leveraging resources from Traditional Chinese Medicine (TCM). Through systematic translation and filtration, we adapted TCM knowledge graphs to reflect TVM’s unique elements, addressing a gap in the documentation of Vietnamese medical knowledge. Our knowledge graph, VieMedKG, serves as a foundational resource for future research and applications in the TVM domain. By providing a curated benchmark for knowledge graph-based question answering, we have laid the groundwork for further developments in QA systems for traditional medicine. Our experiments demonstrate the feasibility and potential of using advanced language models and retrieval-augmented generation techniques in this context.

**A Description of VieMedKG**

